# A *de novo* transcriptomic atlas of early embryo development in the Arabian killifish

**DOI:** 10.64898/2026.07.17.738637

**Authors:** Ruth Y. Akinmusola, Rashid Minhas, Paul O’Neill, Koto Kon-Nanjo, Tetsuo Kon, Yasuhito Shimada, Mark Ramsdale, Tetsuhiro Kudoh

## Abstract

The Arabian killifish (*Aphaniops dispar*) is new tractable vertebrate model system for developmental, ecological and biomedical research, including drug screening, pharmacological and infection biology studies. It is a relatively small euryhaline teleost with broad thermal tolerance and adaptability across a wide range of salinities from freshwater to hypersaline habitats. The embryos and early larvae are tolerant to environmental stressors and exhibit a delayed period of nutritional independence before hatching. This advantage offers an extended window for experimenting on the early developmental processes. Here, we describe time-course gene expression profiling of Arabian killifish embryos across nine developmental time points, from the 1-cell stage to the larval pre-hatching stage. Clustering of dynamic expression profiles for 27,564 Trinity genes revealed coordinated transcriptional modules corresponding to the maternal, blastula, maternal-to-zygotic transition (MZT)-related, gastrulation, organogenesis and larval maturation stages. The maternal stage displayed a highly distinct expression profile, dominated by maternal-specific transcripts that are rapidly degraded during the MZT. The later stages, from 48 hpf onward, revealed a shift from early regulatory mechanisms to the expression of organogenesis-related genes. The ZGA stage showed the conserved up-regulation of many zinc finger-associated genes, consistent with zebrafish and other teleost genomes. Overall, embryo development in *A. dispar* is slower than in zebrafish, with equivalent stages occurring several hours later. We propose a delayed onset of zygotic genome activation (ZGA) in the blastula stage, corresponding to 6 hpf in the Arabian killifish. Taken together, this study provides a transcriptomic resource for mining embryo development-related genes in the Arabian killifish.

## Background and Summary

The Arabian killifish (*Aphaniops dispar*) is an emerging vertebrate model with exceptional experimental versatility. It is a relatively small-sized fish belonging to the Cyprinodontoidei, a diverse clade that includes killifishes, pupfishes and topminnows^1^. Members of this group are characterised by pronounced ecological specialisation and remarkable tolerance to environmental stressors^2–4^. Within this clade, *A. dispar* is particularly notable for its natural resilience to extreme fluctuations in temperature, salinity, and UV exposure^5–7^. It also possesses distinctive fluorescent pigment cells (fluoroleucophores) quite detectable at the larval stages^8^. Furthermore, the chorion is highly transparent, facilitating live imaging and reporter assays^8^. The embryos are viable at high temperatures (37^∘^C and 42^∘^C), and this offers a unique advantage of modelling pathogen infection dynamics at a temperature physiologically relevant to humans^9–11^. Combined with a short generation time and high fecundity, these attributes make *A. dispar* a tractable vertebrate system for developmental, ecological and biomedical research, including drug screening and pharmacological studies.

The morphological processes underpinning the transition from fertilised fish egg to free-living larva have been broadly characterised in teleosts^12–15^. Importantly, in *A. dispar,* the developmental stages are well defined, and independent feeding begins at 13 dpf in breeding populations at 28 ^∘^C, enabling researchers to conduct more experiments before the species becomes protected under welfare regulations^16^. However, these morphological descriptions have not yet been integrated with the underlying molecular gene networks or regulatory pathways. Previous transcriptomics studies in teleost models have revealed dynamic shifts in gene expression across early developmental milestones, including maternal transcript utilisation and the maternal-zygotic transition^17–23^. Other studies have conducted broader time-course analyses across multiple embryonic stages^24,25^.

Species-specific embryonic transcriptomes are essential because fish lineages differ markedly in the timing of the maternal-zygotic transition, polyadenylation dynamics, early zygotic activation, somitogenesis, organogenesis, larval physiology, independent feeding and hatching^18,21,24,26,27^. Currently, genomics resources for *A. dispar* are scarce, limiting functional and comparative studies. A time-course *de novo* analysis of relative mRNA expression patterns will provide a temporal atlas of the regulatory genes governing embryo development^24^. It will also enable the detection of maternal transcripts not supported by somatic genome-based mapping. This will enable the identification of the core genes and pathways involved at distinct developmental time points most especially species-specific developmental delays. Furthermore, this will serve as a foundation for a genetic toolkit for downstream applications in ecotoxicology and infection biology in the Arabian killifish.

In this study, we performed a *de novo* RNA-seq time-series analysis across key embryonic stages of the Arabian killifish. The genome-wide gene expression profiling captured major developmental time points, including maternal transcript expression, blastula formation, maternal-zygotic transition, and organogenesis. By resolving clusters of genes expressed at these stages, we identified stage-specific mRNA signatures and established a foundational dataset for comparative studies in vertebrate developmental biology.

## Methods and Results

### Sample collection, RNA extraction and quality metrics

Embryos of *Aphaniops dispar* were collected at 0.5 hpf, 6 hpf, 12 hpf, 24 hpf, 48 hpf, 72 hpf, 5 dpf, 7 dpf and 11 dpf. These nine key stages captured developmental progression from the 1-cell period through the pre-hatching stage^16^ (Figure 1, Table 1). Three biological replicates per stage were flash-frozen in liquid Nitrogen and stored at -80°C. Total RNA was extracted using the Monarch total RNA miniprep kit (New England Biolabs Catalog #T2010). RNA quality was assessed using Qubit fluorometer for concentration, Agilent Bioanalyzer (Agilent Technologies, Inc.) for RNA integrity number (RIN ≥ 7) and NanoDrop spectrophotometer for purity (A260/280 ratio 1.8–2.0).

**Figure 1:**
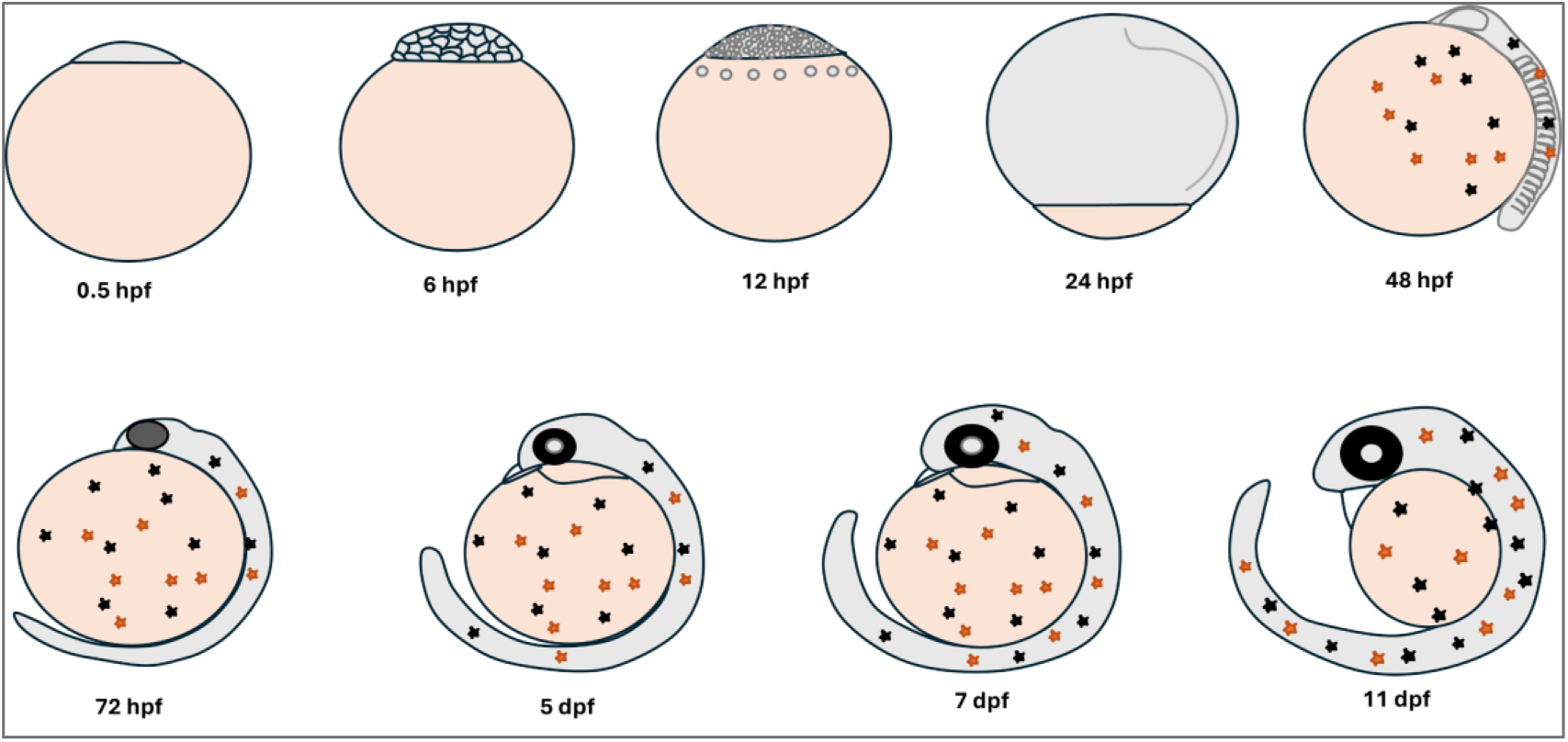
The sampled *A. dispar* developmental stages from 0.5 hpf to 11 dpf. The time records are hours post fertilisation (hpf) and days post fertilisation (dpf). Some of the body parts become visible at around 48 hpf. The black dots represent melanophores while the red dots represent fluoroleucophores.

**Table 1:** Sampled *A. dispar* embryos and their corresponding stages. The numbers in the bracket are developmental staging identifiers where each stage corresponds to specific anatomical or morphological features in the Arabian killifish embryos^16^.

| Sample | Stage (*A. dispar*) |
| --- | --- |
| Maternal (0.5 hpf) | 1-cell (stage 1) |
| 6 hpf | Early to mid-blastula (stages 5-7) |
| 12 hpf | Late blastula (stages 12-13) |
| 24 hpf | 90% epiboly (stage 24) |
| 48 hpf | Early organogenesis (stage 48) |
| 72 hpf | Mid organogenesis (stage 72) |
| 5 dpf | Early larval/late organogenesis (stage 120) |
| 7 dpf | Developing larval stage (stage 144) |
| 11 dpf | Late larval/pre-hatching (stage 264) |

### Library preparation and sequencing

RNA-seq libraries were prepared using the Illumina® Stranded mRNA prep kit according to the manufacturer’s protocol. Briefly, poly(A)+ RNA was extracted and fragmented prior to cDNA synthesis with the second strand marked for later degradation. Libraries were indexed, amplified and purified before quality assessment using the Agilent Bioanalyzer (Agilent Technologies, Inc.) and Qubit fluorometer. Sequencing was performed on an Illumina NovaSeq 6000 platform, S1 flowcell, generating a total of 874,900,014 raw reads from 150 bp paired-end libraries. Low quality (<Q22) and adapter sequences were removed from the ends of the reads with Fastp v0.23.4 with default parameters. The quality of the raw and clean sequencing reads was assessed using FastQC v0.11.9^28^. These quality metrics were aggregated and summarised with MultiQC v1.16^29^ for all samples (Table S1A). All the samples exhibited high mean Phred quality scores (>Q30), high quality per sequence quality scores (denoted as a sharp peak at >Q30), consistent GC content distribution and a negligible per base N content at all base positions (Figure 2, Table S1A).

**Figure 2:**
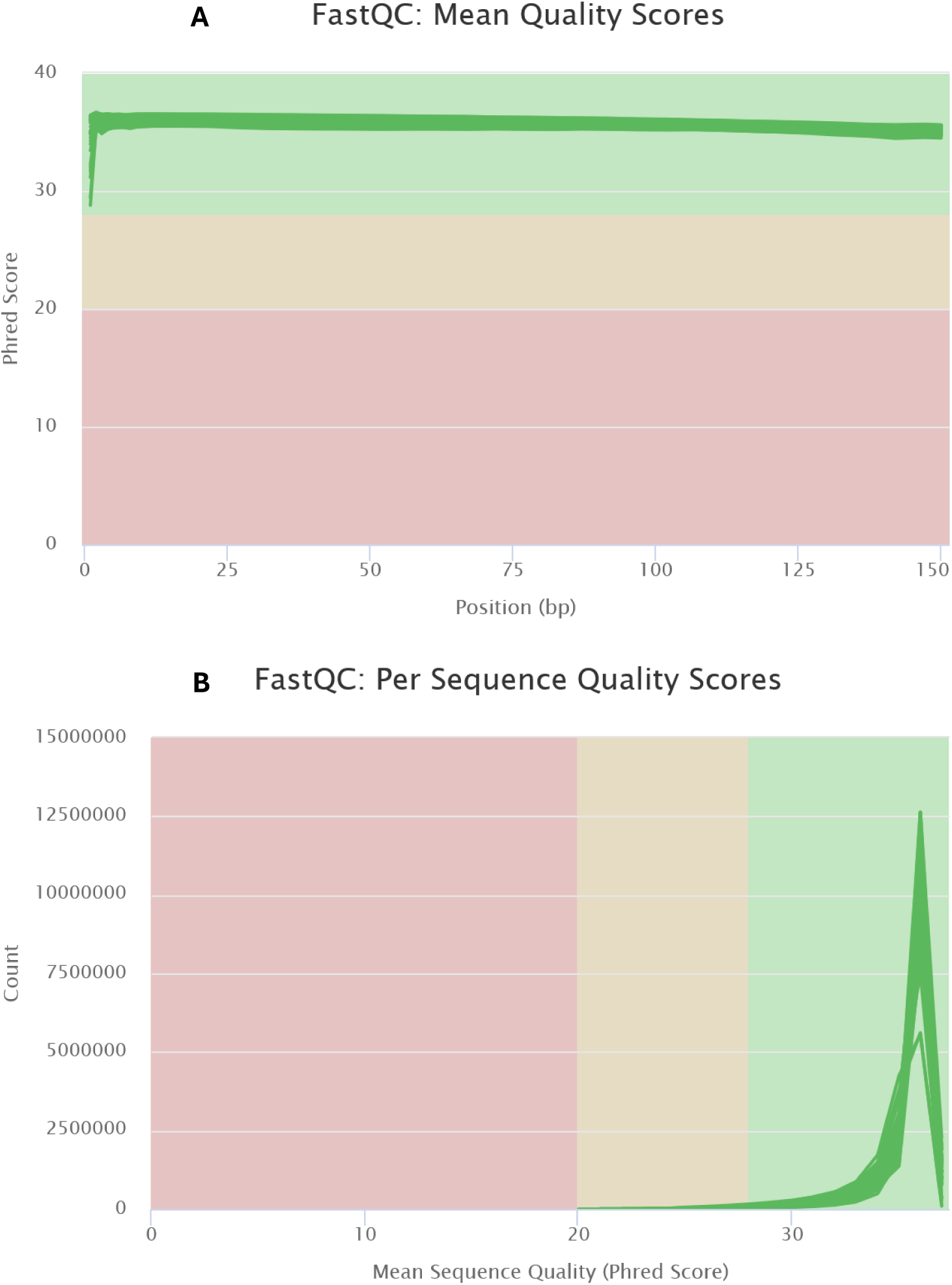
MultiQC assessment of the trimmed sequencing reads from all chosen developmental stages (maternal to hatching). (**A**) Mean quality scores across all base positions. (**B**) Per sequence quality scores across all Phred scores.

### Assembly completeness and mapping rates

Gene completeness of the assembly was evaluated with BUSCO completeness scores of 95% and 88% for Actinopterygii and Cyprinodontiformes lineages datasets, respectively (Figure 3A). This indicates a strong representation of conserved genes across fish lineages. Trimmed reads from each biological replicate and developmental stage were aligned back to the *de novo* transcriptome with HISAT2 v2.2.1^29^ in a paired-end mode. Alignment rates ranged from 86% to 95%, demonstrating a high-quality representative Trinity assembly (Figure 3B).

**Figure 3:**
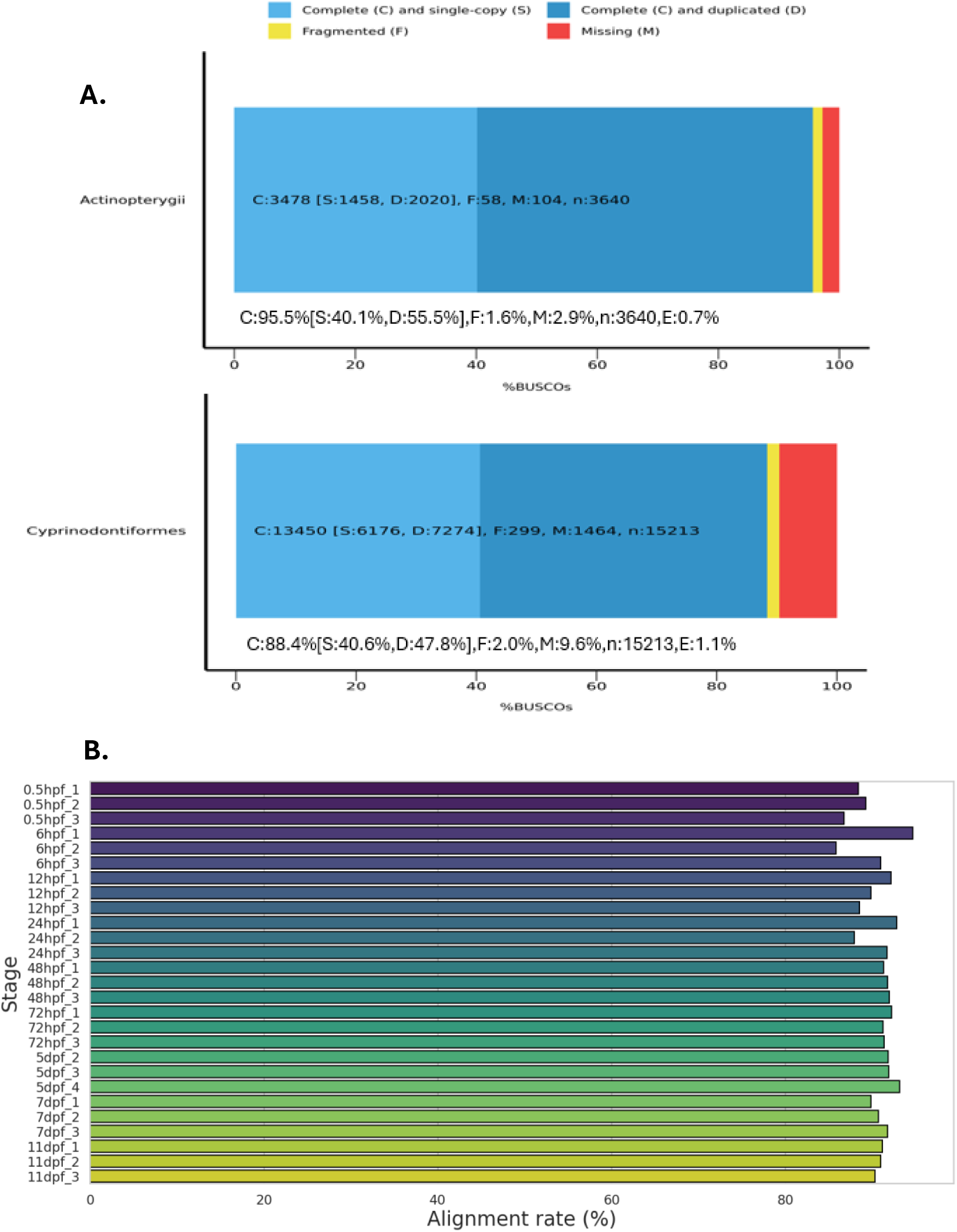
Assembly completeness of the *de novo* transcriptome. (**A**) BUSCO assessment of gene completeness. The results were classified on the proportion of complete (single-copy (S) or duplicated (D)), fragmented (F) or missing (M) BUSCOs. (**B**) Mapping rates of the individual embryo stages against the *de novo* transcriptome.

### De novo transcriptome assembly and redundancy reduction

All clean reads were subjected to SortMeRNA^30^ to remove ribosomal RNA. The reads were combined and assembled *de novo* using Trinity^31^ with default parameters and i*n silico* read normalisation enabled. The TrinityStats.pl utility and TransRate were used for assembly statistics and length distribution^31,32^. Trinity reconstructed transcript sequences representing alternative isoforms and gene clusters from a total of 1,015,470 genes with a combined length of 598,733,144 bp (Table S1B). The GC content of the assembly is 47.11% (Table S1B), within the range reported for a salinity-based *de novo* transcriptome of the Arabian killifish^33^. The initial Trinity assembly contained many contigs less than 200 bp indicating the presence of many short transcripts. (Table S1C). This relatively high number of short contigs which likely reflects transcript isoform diversity and fragmentation of low-abundance transcripts, consistent with the expectations from a *de novo* transcriptome assembly^34,35^.

Downstream analyses were restricted to predicted protein-coding transcripts identified using TransDecoder (https://github.com/TransDecoder/TransDecoder/wiki v5.5.0. TransDecoder identified the longest and highest-confidence open reading frames (ORFs) within the Trinity transcripts by integrating coding potential, ORF length, and homology evidence from Pfam domain searches. The predicted protein-coding transcripts were clustered at a 90% identity threshold using cd-hit v4.8.1^36^ (cd-hit-est) to reduce redundancy, resulting in a mean ORF prediction rate of 99.84% (Table S1C). This reduced the overall mapping rate and total number of genes from 1,015,470 to 149,382 with increased average contig length (695.73 bp) and GC content (52.6%) (Table S1B-D). The BUSCO assessment of clustered transcriptome sequences showed a reduction in duplicated genes from 55.5% to 6.3%, while maintaining a high gene completeness of 94% in the Actinopterygii_odb10 lineage (Table S1D). Thus, the clustered dataset represents a reasonable degree of non-redundancy and an acceptable coverage of protein-coding transcripts for downstream analysis.

### Functional annotation

The assembled transcripts were functionally annotated with BLASTX and BLASTP searches against the Uniprot/Swiss-Prot database following Trinotate^37^ default parameters. Searches were conducted with an E-value cutoff of 1e-5, a maximum of one target sequence per query (max_target_seqs 1) and the results were reported in tabular format (outfmt 6). Gene ontology (GO) terms including biological process, molecular function and cellular component, and biological trait associations (developmental, induction and tissue specificity) were assigned using the Uniprot ID mapping tool.

Open-reading frames were predicted from the non-redundant transcripts using TransDecoder (https://github.com/TransDecoder/TransDecoder/wiki-v5.5.0) with default parameters, retaining the longest ORF per transcript. Homology support was enabled by incorporating Pfam domain searches via HMMER v3.3.2^38^ and BLASTP hits against the Swiss-Prot database to improve ORF prediction. The TransDecoder-predicted protein sequences were functionally annotated using the eggNOG-mapper v2^39^ with the eggNOG 5.0 database, employing DIAMOND in sensitive (-m diamond) and including all gene ontology evidence (go_evidence all). All gene to transcript annotations were integrated and visualised with Trinotate v4.0.2^37^, incorporating the top BLASTX hits against the Swiss-Prot database along with BLASTP, Pfam and eggNOG for a comprehensive annotation report. A total of 89,643 Trinity genes were annotated resulting in an overall annotation rate of 60.01%.

#### Differential expression

Transcript abundance estimates were generated using Salmon v1.10.3^40^, which produced quant.sf files containing TPM values, NumReads (estimated) counts and effective lengths. Gene-level count data were analysed using DESeq2 v1.34.0^41^. Transcript to gene-level abundance estimates generated by Salmon were imported into DESeq2 using the tximport package^42^. A DESeq2 dataset was constructed from the tximport object using the experimental design formula ∼stage, where stage represents developmental time points. Size factor normalisation was applied across all samples to account for library size differences. Normalised gene-level expression values were extracted with DESeq2 counts (normalised = TRUE) (Figure 4A, Table S2). For gene-level summaries, only the top BLASTX hit was retained as it provided the richest annotation (Figure 4B). Pairwise contrasts between developmental stages were evaluated with the results() function. Significant DEGs were identified using thresholds of log2FC ≥ 1, FDR-adjusted *p*-value < 0.05, and an average normalised count of ≥ 5 in at least two stages (Table S2). Volcano and bar plots were generated with ggplot2, dplyr, and ggrepel for label positioning^43,44^. The highest overlap with the DESeq2-filtered genes occurred at TPM ≥ 0.5 in ≥ 2 stages, indicating the recovery of all genes with approximately a TPM threshold of 0.5 (Table S3).

**Figure 4:**
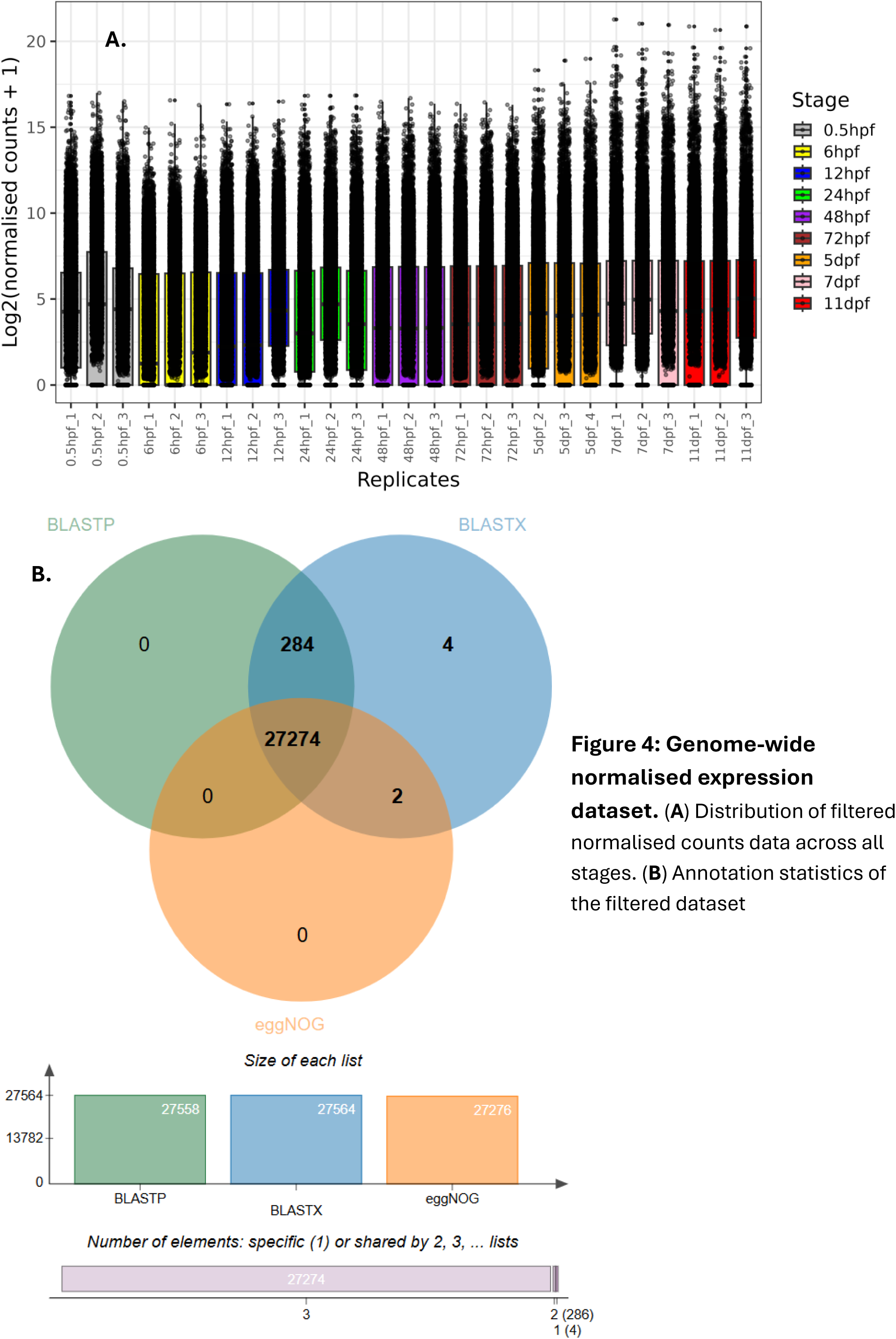
Genome-wide normalised expression dataset. (**A**) Distribution of filtered normalised counts data across all stages. (**B**) Annotation statistics of the filtered dataset

### Clustering relationships across the developmental stages revealed major developmental transitions

#### Expression similarity is stage-specific

Complex heatmap in R was used for clustering patterns of differential expression across all stages^45^. Pairwise relationships in the data across all stages were visualised with RNAlysis v4.2.0^46^. Hierarchical clustering grouped the genes based on stage-specific expression profiles and captured genome-wide co-regulated gene transcriptional patterns (Figure 5D). The maternal stage exhibited the most dissimilar expression profile to all the other developmental stages (Figure 5A and D). This pattern reflects the detection of maternal-specific and maternally-supplied transcripts that undergo degradation following the maternal-to-zygotic transition (MZT). In addition, a volcano plot showed the top DEGs (differentially expressed genes) between the 0.5 hpf and 11 dpf stages (Figure 5B). Genes such as *dnmt1* (*DNA (cytosine-5-)-methyltransferase 1*) show strong maternal enrichment but are significantly downregulated by 11dpf (Figure 5C). This pattern is consistent with their roles in early embryonic regulation. *dnmt1* is essential for maintaining the inherited maternal methylome during the earliest cell cycles, thereby ensuring proper epiboly and preventing embryonic lethality around gastrulation^47–49^. In contrast, we observed the differential up-regulation of *idh3a* (*isocitrate dehydrogenase (NAD^+^) 3 catalytic subunit alpha*)*, slc25a11* (*solute carrier family 25 member 11*) and *hsp70* (*heat shock protein 70*) at 11dpf (Figure 5C).

**Figure 5:**
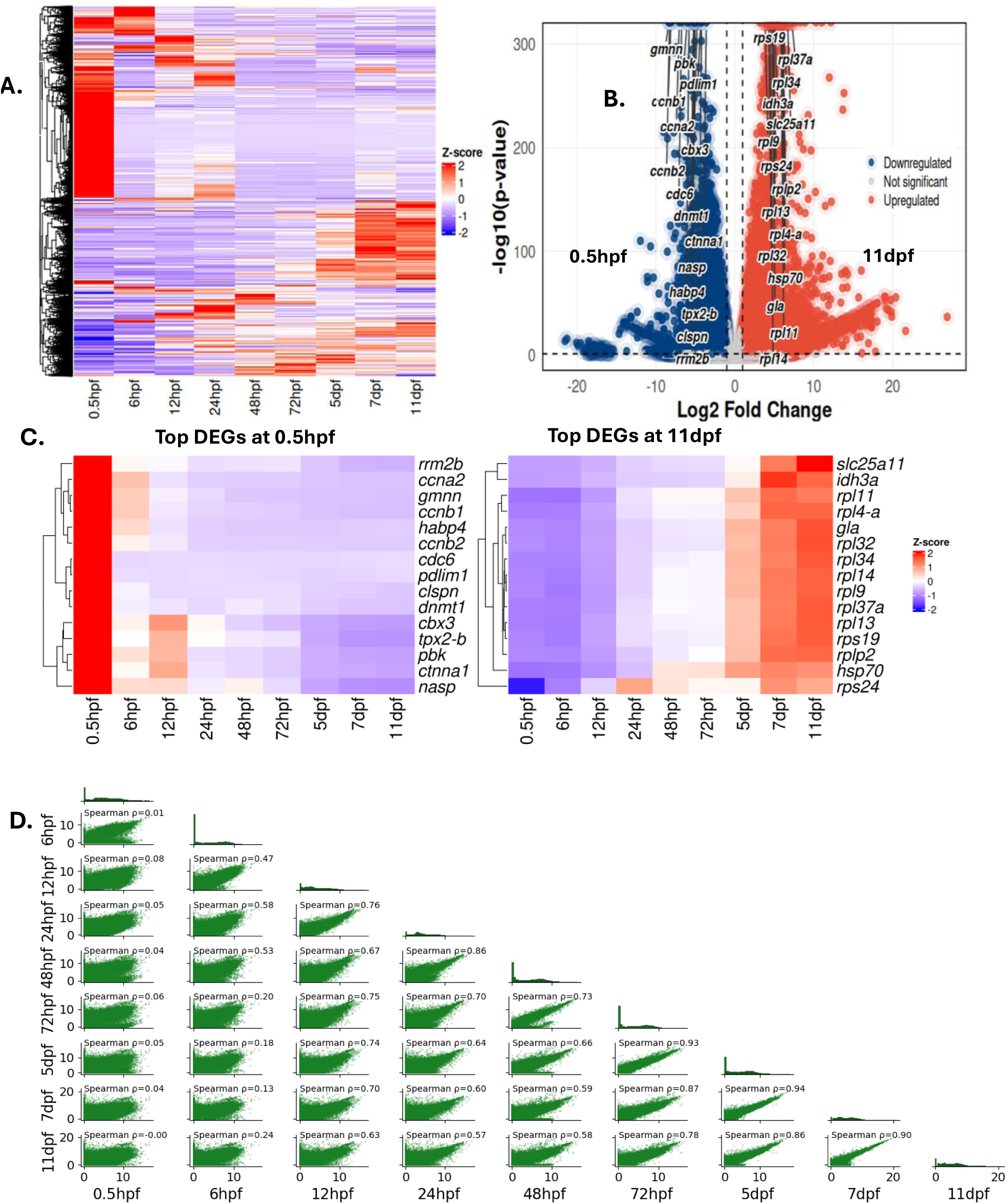
Genome-wide gene expression patterns across all the selected stages. (**A**) Clustering patterns of size-factor normalised counts of the 27,564 filtered genes across all the selected developmental stages. Red Z-scores indicate high relative expression, white Z-scores show average expression and blue Z-cores indicate low relative expression. (**B**) A volcano plot showing the top DEGs between the maternal stage and 11dpf. (**C**) Heatmaps showing expression patterns of the top DEGs at 0.5hpf and 11dpf. (**D**) A logarithmic scale pairplot showing relationships between samples across all the embryo stages. *ρ* represents Spearman correlation coefficient between each pair of embryo stages.

The 6–12 hpf interval (early to late blastula) represents a transitional developmental cluster, showing moderate correlation with the shift from continuous cleavage to the formation of the yolk syncytial layer (YSL) prior to gastrulation^16^ (Figure 5A and D). The 24-72 hpf clusters are clearly distinguishable from both earlier and later developmental stages, indicating transcriptional up-regulation associated with epiboly, gastrulation, axis formation, and the onset of early organogenesis^16,50^ (Figure 5A and D). Stages 24-72 hpf exhibit some interstage correlation, indicating the presence of shared gene expression (Figure 5D). The positive correlation in this expression window may be explained by a continuous developmental trajectory with highly synchronised biological transitions for the first major organs to be formed^16^. Notwithstanding, the highest levels of positive correlation were observed between the late developmental stages from 5 dpf (Figure 5A and D). These 5-11 dpf clusters reveal a late-stage gene expression signature, consistent with late organogenesis and the establishment of larval physiology required for hatching^16^.

#### Mfuzz soft clustering identified distinct temporal gene expression patterns across developmental stages

To identify distinct temporal gene expression patterns across the developmental stages, soft clustering was performed using the Mfuzz package v2.54.0, R/Bioconductor. The normalised expression matrix was converted into an ExpressionSet object and standardised prior to clustering. The fuzzifier m was estimated with mestimate(), and the number of optimum clusters (c) *was* selected based on the Dmin function and biological relevance^51^. This analysis revealed nine distinct expression clusters, with cluster 9 containing the largest number of genes (Figure 6, 7A and B, Table S8A-J). The nine clusters revealed diverse developmental time points and the prominent/mid-timepoint in the up-regulation peak was used to assign cluster names (Figure 6).

**Figure 6:**
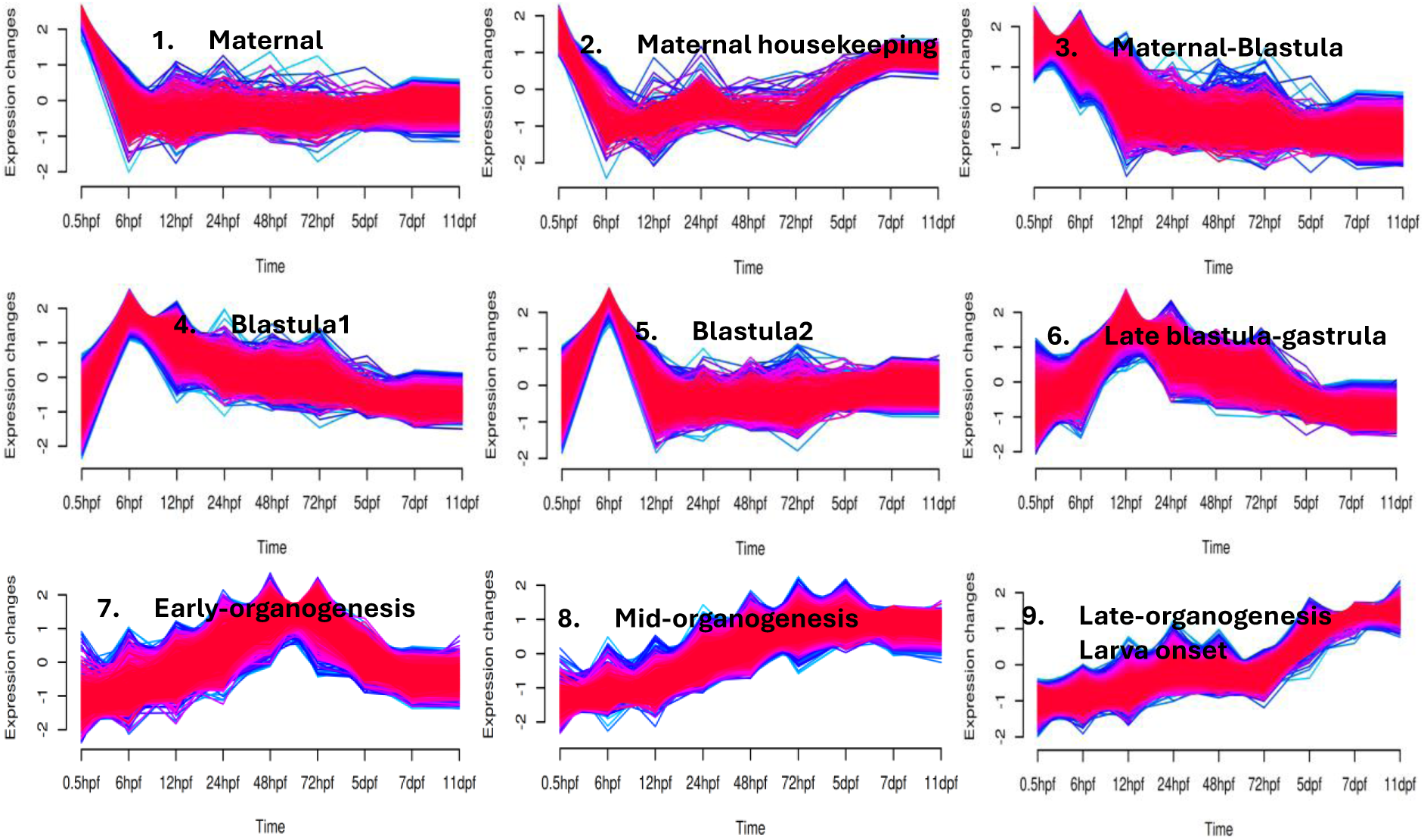
Time series of gene expression trends across all the developmental stages from maternal to 11 dpf. Transcriptome-wide clustering with Mfuzz showed nine distinct patterns of gene expression from 0.5 hpf to 11 dpf.

In the maternal cluster (cluster 1), we detected classic maternal genes that have been studied in Zebrafish such as *dazl* (*deleted in azoospermia-like*), *bol* (*boule homolog, RNA-binding protein*), and maternally supplied transcripts in the early cleavage stages such as *hwa* (*huluwa)* (Figure 8). Cluster 2 corresponds to maternal housekeeping genes. Cluster 3 contains genes highly expressed between the maternal to blastula stages such as *ccne1* (*cyclin E1*) and *nanog* (*nanog homeobox*). Two blastula clusters were detected (cluster 4 and 5) with cluster 4 containing classic blastula genes well studied in teleosts. These include early to mid-blastula genes such as *pou5F1* (*POU domain, class 5, transcription factor 3*)*, pum2* (*Pumilio 2, conserved RNA-binding protein*) and *chaf1a* (*chromatin assembly factor 1*) (Figure 8). We also observed a cohort of rapidly induced Zinc-finger domain associated genes within cluster 4 and 5, which corresponds to the early to mid-blastula stage (Figure 9, Tables S6 and S7). These reflected the ZGA-associated Zinc finger gene expansions observed in other teleosts such as zebrafish and *Cynoglossus semilaevis*^23,24^. Zn-finger TFs are typically among the first classes of transcription factors activated at ZGA^52,53^. A substantial proportion of them are characterised by the presence of C2H2 Zinc finger domains (Figure 9, Table S6B and S7B). The top significant gene ontology (GO) terms in the Zinc-finger associated clusters are mainly involved in metabolic processes, nuclear processes and DNA binding transcription factor activity (Figure S2-3). These significant GO terms are consistent with established roles of Zinc finger proteins in transcriptional and post-transcriptional gene regulation^54^.

The maternal (cluster 1), maternal-blastula (cluster 3) and blastula 1 (cluster 4) clusters showed overlaps in gene membership (Figure S1). GO terms associated with cell division are significantly enriched in cluster 3, 4 and 6 which correspond to early embryogenesis stages characterised by rapid developmental progression (Figure 7C). Notably, the stages covered by these clusters occur within the first 24 hours of development in the Arabian killifish. Cluster 4 (blastula 1) exhibited the highest enrichment in top significant GO terms associated with DNA damage response and regulation of transcription by RNA polymerase II (Figure 7C). Among them all, the largest cluster (late organogenesis/larva onset (cluster 9)) contains the highest number of UniProtKB-annotated developmental, induction and tissue specific genes indicating the activation of highly specialised organ-specific transcription (Figure 7A-B). This cluster contains candidate genes required for autonomous life such as *fabp2* (*fatty acid binding protein 2*) a biomarker of gut maturation involved in the activation of lipid uptake^55^ (Figure 8).

**Figure 7:**
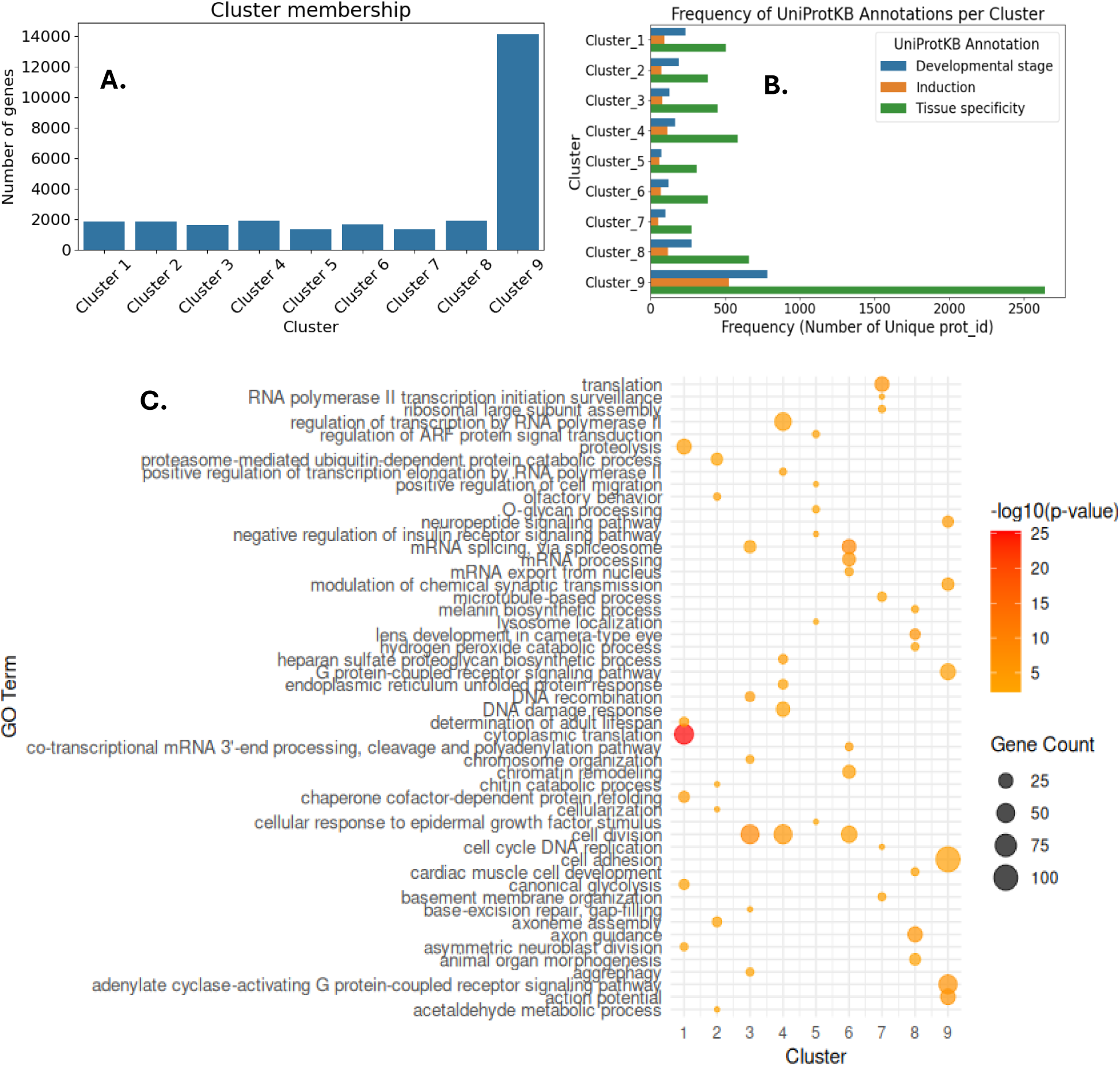
C**l**uster **membership and GO analysis of the candidate genes in all the developmental stages from maternal to 11 dpf.** (**A**) Total number of Trinity genes in each Mfuzz cluster (**B)** Proportion of genes associated with developmental stage, induction and tissue specificity annotations across the clusters mapped to the UniProtKB database. (**C)** Top significantly enriched GO terms in each cluster. Cluster membership: Cluster 1 (Maternal), Cluster 2 (Maternal housekeeping), Cluster 3 (Maternal-Blastula), Cluster 4 (Blastula 1), Cluster 5 (Blastula 2), Cluster 6 (Late blastula-gastrula), Cluster 7 (Early organogenesis), Cluster 8 (mid-organogenesis), Cluster 9 (late organogenesis/larva onset). The GO terms are biological processes.

**Figure 8:**
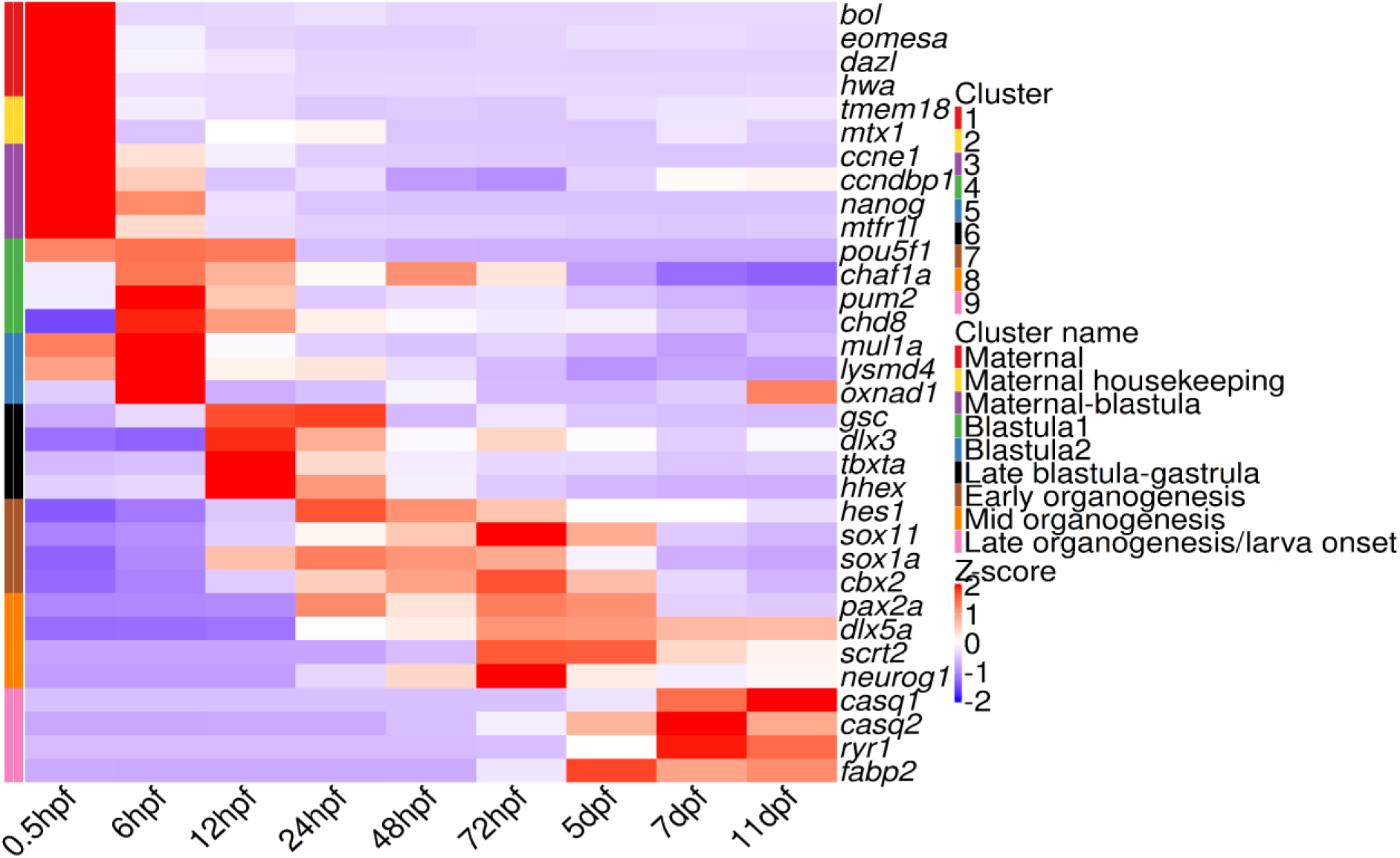
Expression patterns of selected embryonic developmental genes enriched in the Mfuzz clusters. The deseq2 size factor normalised counts for each gene were scaled with z-scores prior plotting. The red z-scores show upregulation while blue represent downregulation. The colour palettes on the left represent the Mfuzz cluster number and cluster names.

### Technical validation

Complementary validation strategies were employed to assess the quality and completeness of the *de novo* transcriptome assembly. These include read representation, gene completeness, redundancy assessment metrics (Table S1A-E). The representative sequences clustered with cd-hit captured the quantitative signal of the original assembly without a significant shrinkage of the core BUSCO genes (Table S1E). We compared gene expression using gene level counts derived from DESeq2 size factor normalised counts (Table S2), Salmon’s transcript-level quasi-mapping quantification (Table S4) as well as transcript per million (TPM) estimates (Table S5). Gene expression estimates were highly consistent across different quantification methods (Figure 9, Tables S4 and S5) for the Trinity genes. Correlations with Salmon gene counts were slightly higher than with TPM, indicating the reliance of DESEQ2 and salmon on count measures (Figure 9). The gene length correction incorporated by TPM resulted in slightly lower correlation but strong agreement across all stages. Furthermore, the principal component analysis revealed the distinct clustering of the maternal stage from all the other stages. Additionally, the early (6hpf, 12hpf and 24hpf) and the late stages (5dpf to 11dpf) revealed distinct transcriptomic landscapes (Figure S4). Furthermore, we also validated the dataset with the expression of known developmental markers in fish embryonic transcriptome datasets such as *dazl* in the maternal stage and *pou5f1* in the blastula stage (Figure 8).

**Figure 9:**
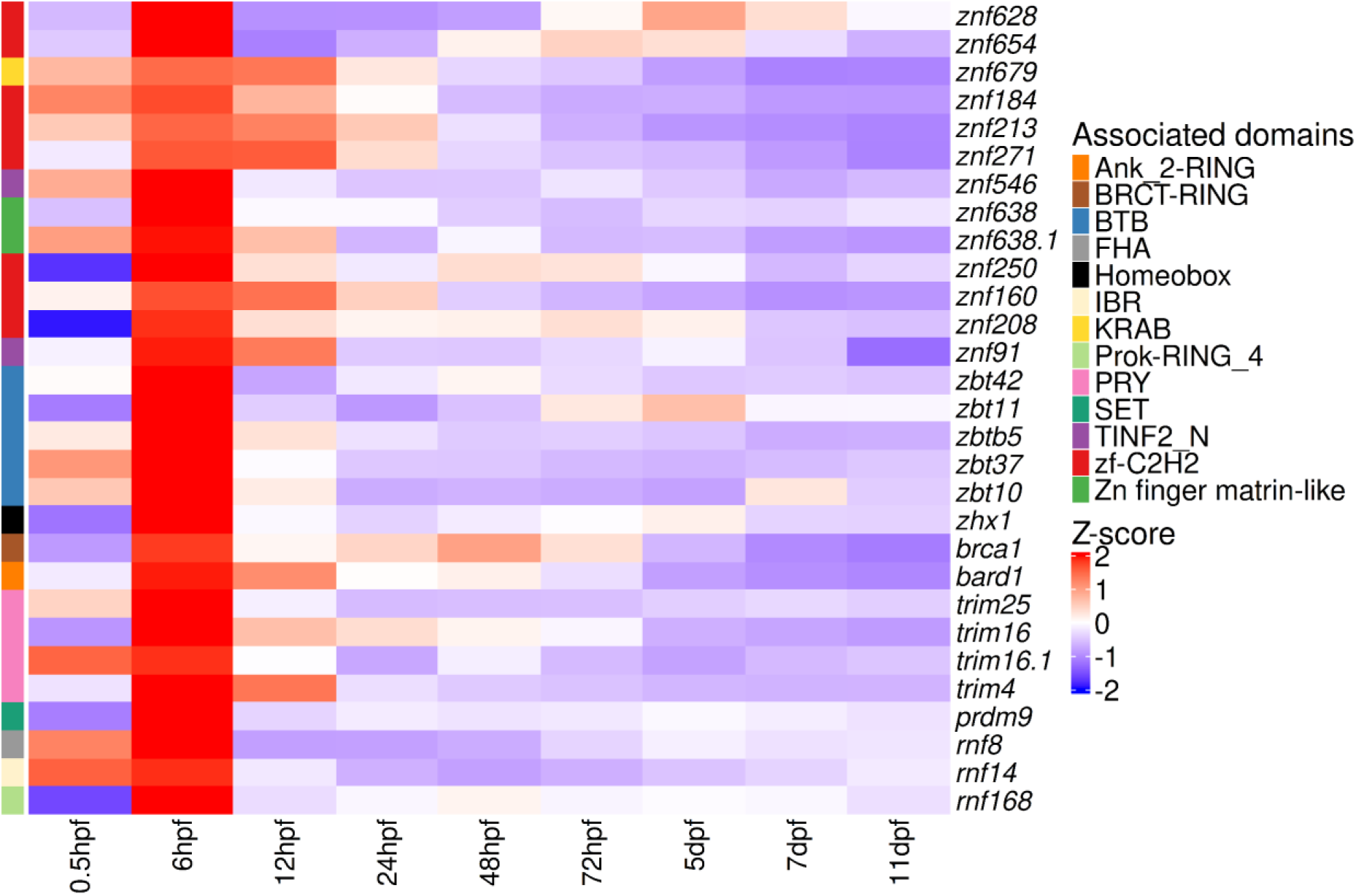
ZGA-related upregulation and the distribution of Zinc finger domain associated genes in the blastula cluster 4. The upper panel shows the expression patterns of Zn-F associated genes across different embryo stages. The bottom panel indicates the functional domains annotated using PFAM. The DESeq2 size-factor normalised counts for each gene were scaled using Z-scores. Ank_2-RING: Ankyrin repeat plus RING domain. BRCT: BRCT domain plus RING domain. BTB: Broad-Complex, Tramtrack, and Bric-à-Brac domain also referred to as POZ domain. FHA: Forkhead-Associated domain. Homeobox: Homeobox DNA binding domain. IBR: In-Between-RING domain. KRAP: Krüppel associated box. Prok-RING_4: Prokaryotic RING-like domain (family 4). PRY: PRY domain. SET: Su(var)3-9, Enhancer of zeste, Trithorax domain. TINF2_N: N-terminal domain of TINF2_N (TRF1 interacting factor 2). zf-C2H2: Cys2-His2 zinc finger. Zn finger matrin-like: Matrin-like Zinc finger domain.

**Figure 10:**
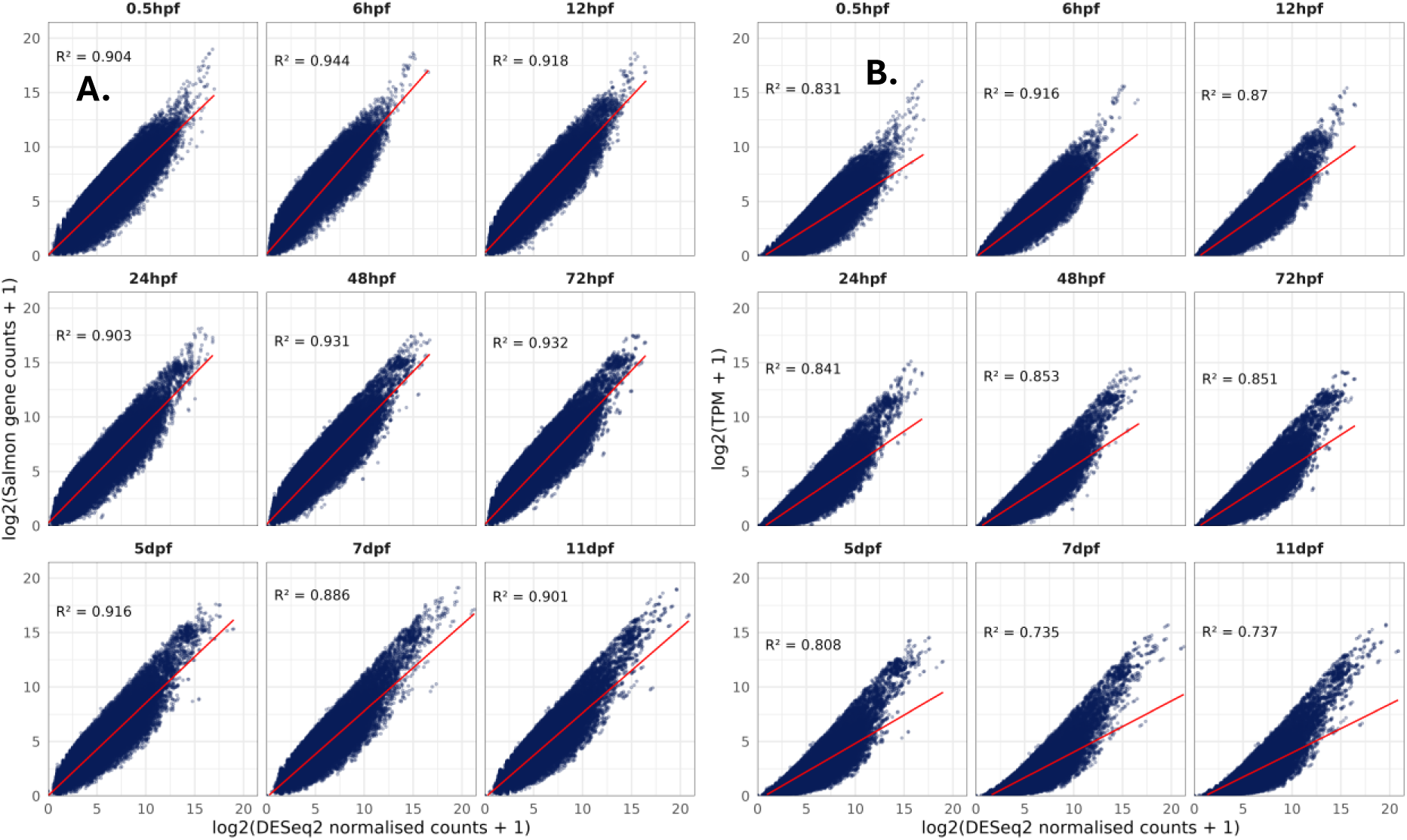
Agreement between expression quantification methods across the developmental stages. (**A**) DESeq2 size factor normalised counts versus salmon gene counts. (**B**) DESeq2 size factor normalised counts versus TPM. The R^2^ values in the legend are stage-specific correlation coefficients.

## Code availability

No custom codes were generated for this study. All analyses were performed using publicly available software following the authors’ recommended manuals and protocols. When specific parameter settings were not explicitly stated, the tools were executed using their default configurations as documented by the software providers.

## Data records

All raw sequencing reads were deposited in the ArrayExpress database under accession number E-MTAB-16987, with the raw sequencing reads available via the European Nucleotide Archive. The corresponding sequencing information, assembly statistics, gene completeness, annotation hits with differential expression dataset and TPM values are available in the supplementary information. The associated trinity assembly, non-redundant assembly, gene and transcript quantification with salmon and annotation files have been deposited in Figshare^56^

## Acknowledgements

The authors acknowledge the funding from NC3Rs (National Centre for the Replacement, Refinement and Reduction of Animals in Research) grant number NC/X001121/1 awarded to Tetsuhiro Kudoh.

## Supplementary Figures

Figure S1: An upset plot showing cluster memberships and overlaps of gene members in the Mfuzz clusters.

Figure S2: Top enriched gene ontologies in Zinc-finger associated genes in cluster 4.

Figure S3: Top enriched gene ontologies in Zinc-finger associated genes in cluster 5.

Figure S4: Principal component analysis of DESeq2 normalisation versus TPM across developmental timepoints.

## Supplementary Tables

Table S1A: Sequencing information

Table S1B: Trinity assembly statistics

Table S1C: Transrate metrics

Table S1D: Read Representation

Table S1E: BUSCO metrics

Table S2: Differential expression statistics and GO terms of all the filtered trinity genes

Table S3: TPM threshold overlaps with the DESeq2-filtered genes

Table S4: Salmon gene count matrix and GO terms of all the filtered trinity genes

Table S5: TPM metrics and GO terms of all the trinity filtered genes

Table S6A: PFAM domains of the Zinc-finger associated genes in cluster 4

Table S6B: gProfiler GO enrichment analysis of the Zinc-finger associated genes in cluster 4

Table S7A: PFAM domains of the Zinc-finger associated genes in cluster 5

Table S7B: gProfiler GO enrichment analysis of the Zinc-finger associated genes in cluster 5

Table S8A: GO annotations of the genes in all Mfuzz clusters based on unique membership.

TableS8B: Mfuzz cluster 1 members

TableS8C: Mfuzz cluster 2 members TableS8D: Mfuzz cluster 3 members

TableS8E: Mfuzz cluster 4 members TableS8F: Mfuzz cluster 5 members

TableS8G: Mfuzz cluster 6 members TableS8H: Mfuzz cluster 7 members

TableS8I: Mfuzz cluster 8 members TableS8J: Mfuzz cluster 9 members

