## Supplemental Figures for "A *de novo* transcriptomic atlas of early embryo development in the Arabian killifish"

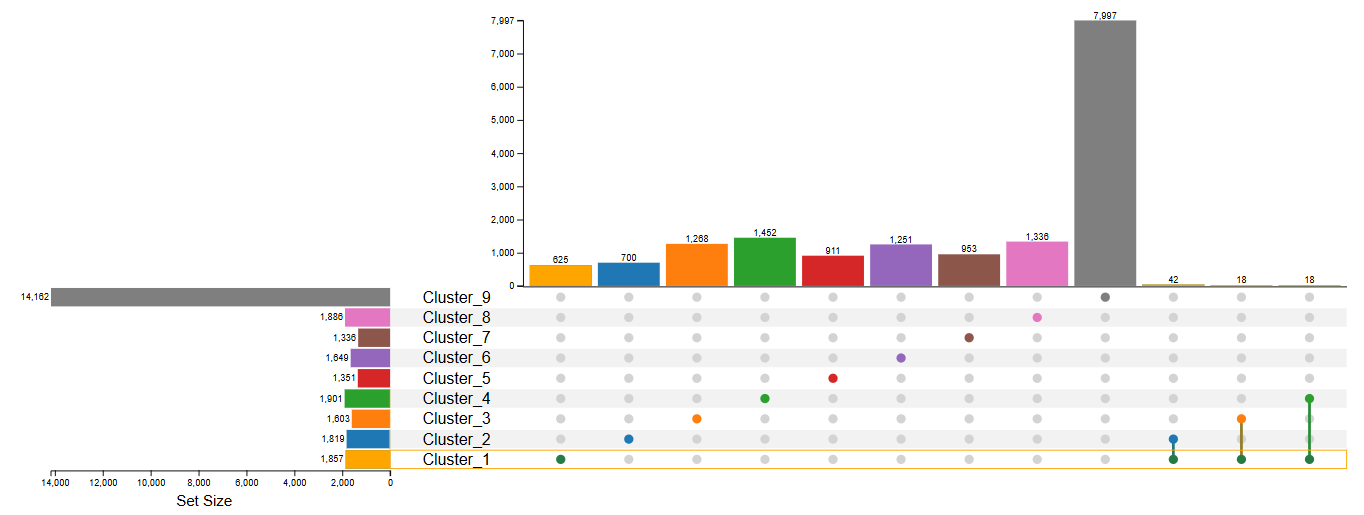
 **Figure S1: An upset plot showing cluster memberships and overlaps of gene members in the Mfuzz clusters.** The first 9 bar plots show the number of distinct genes in each cluster while the last 3 show the number of shared genes.

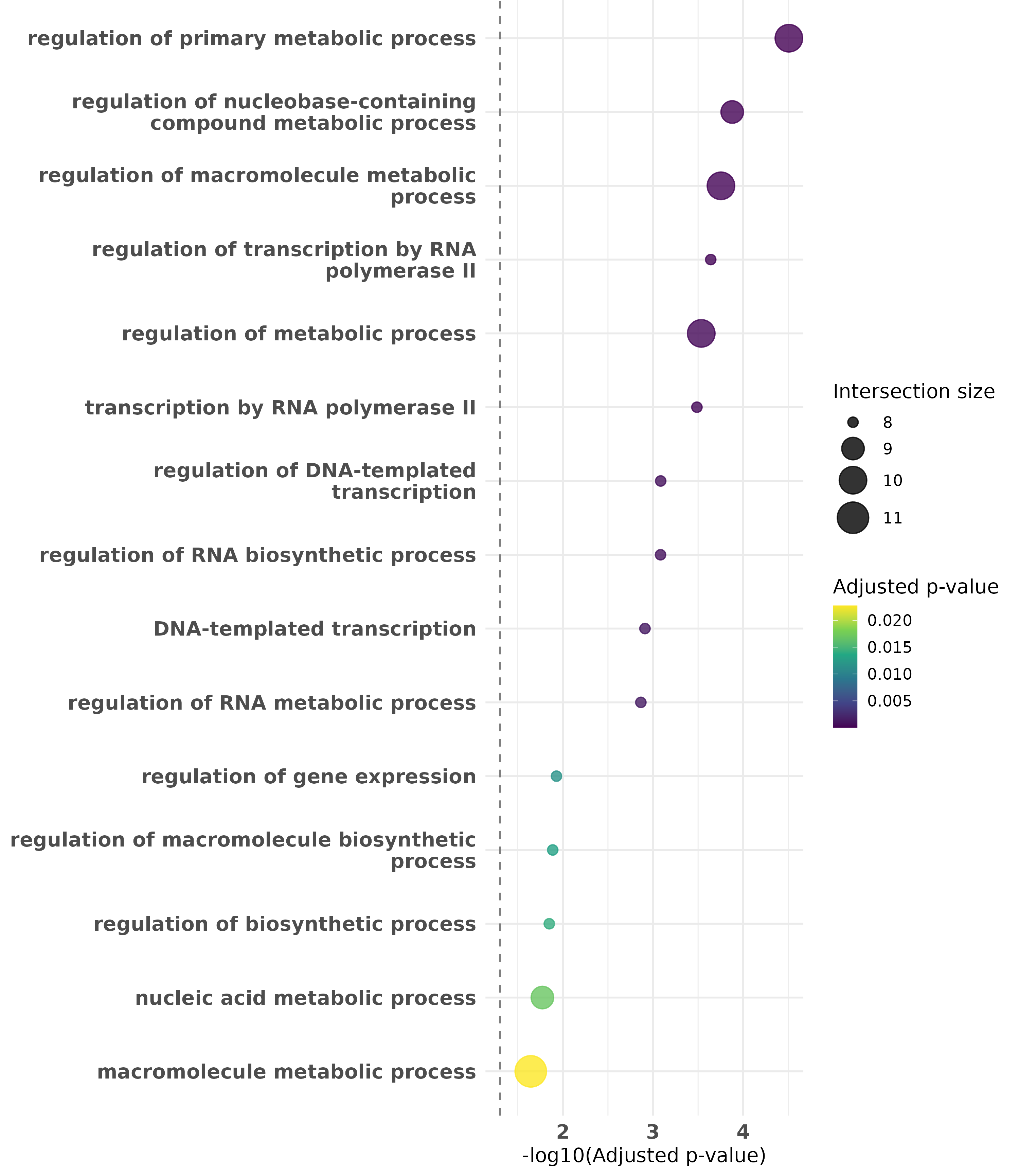

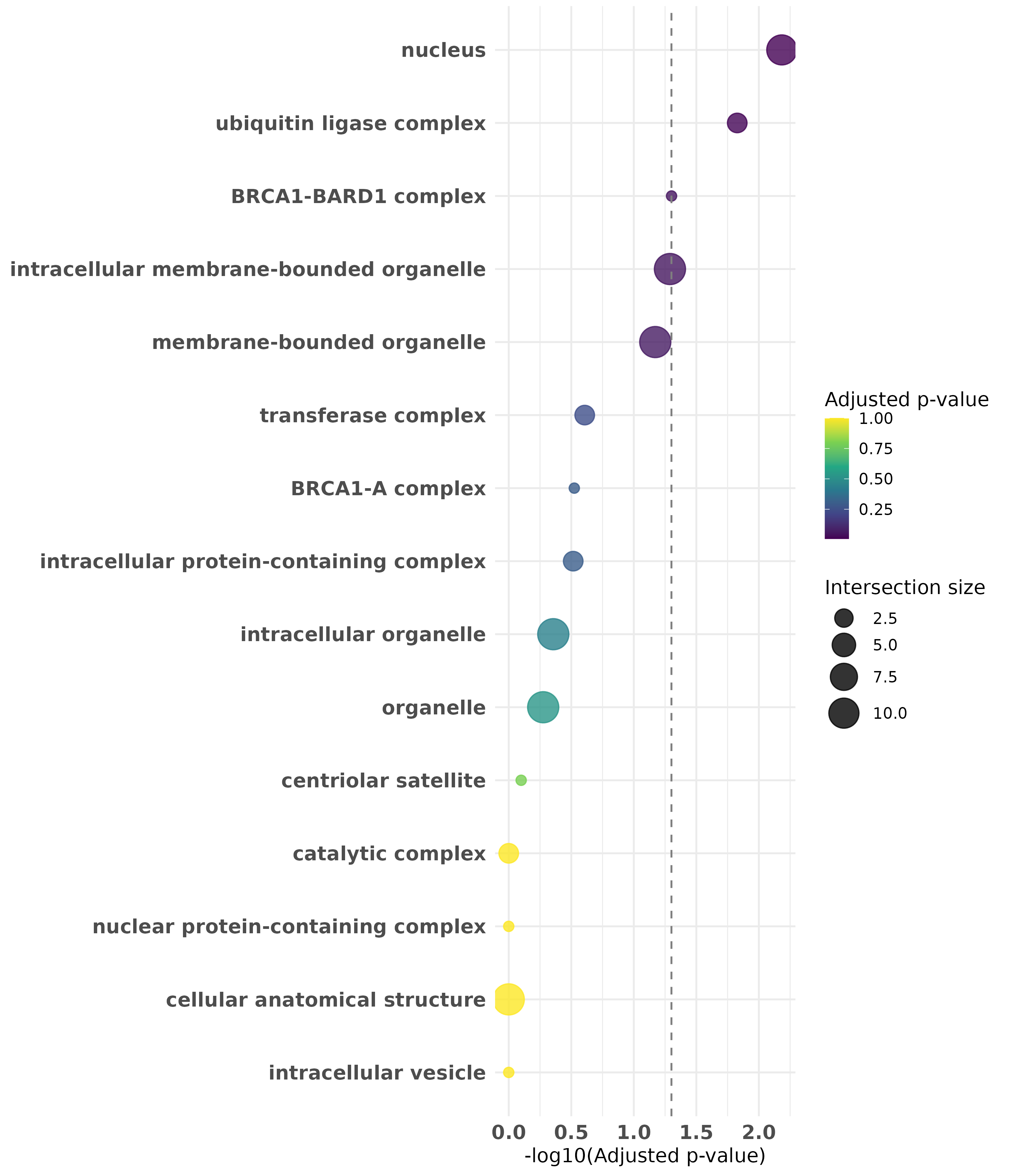

**B**

**A**

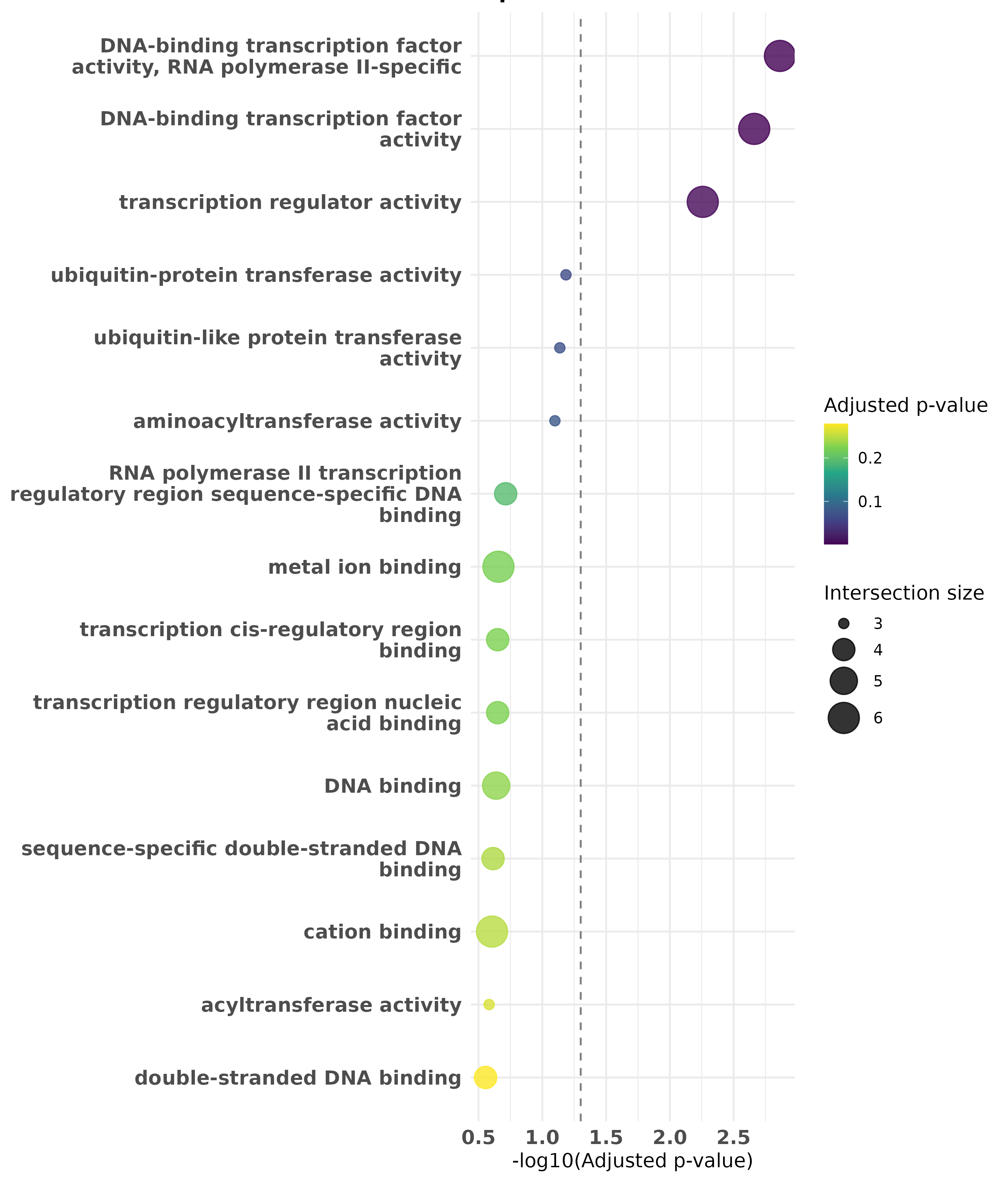

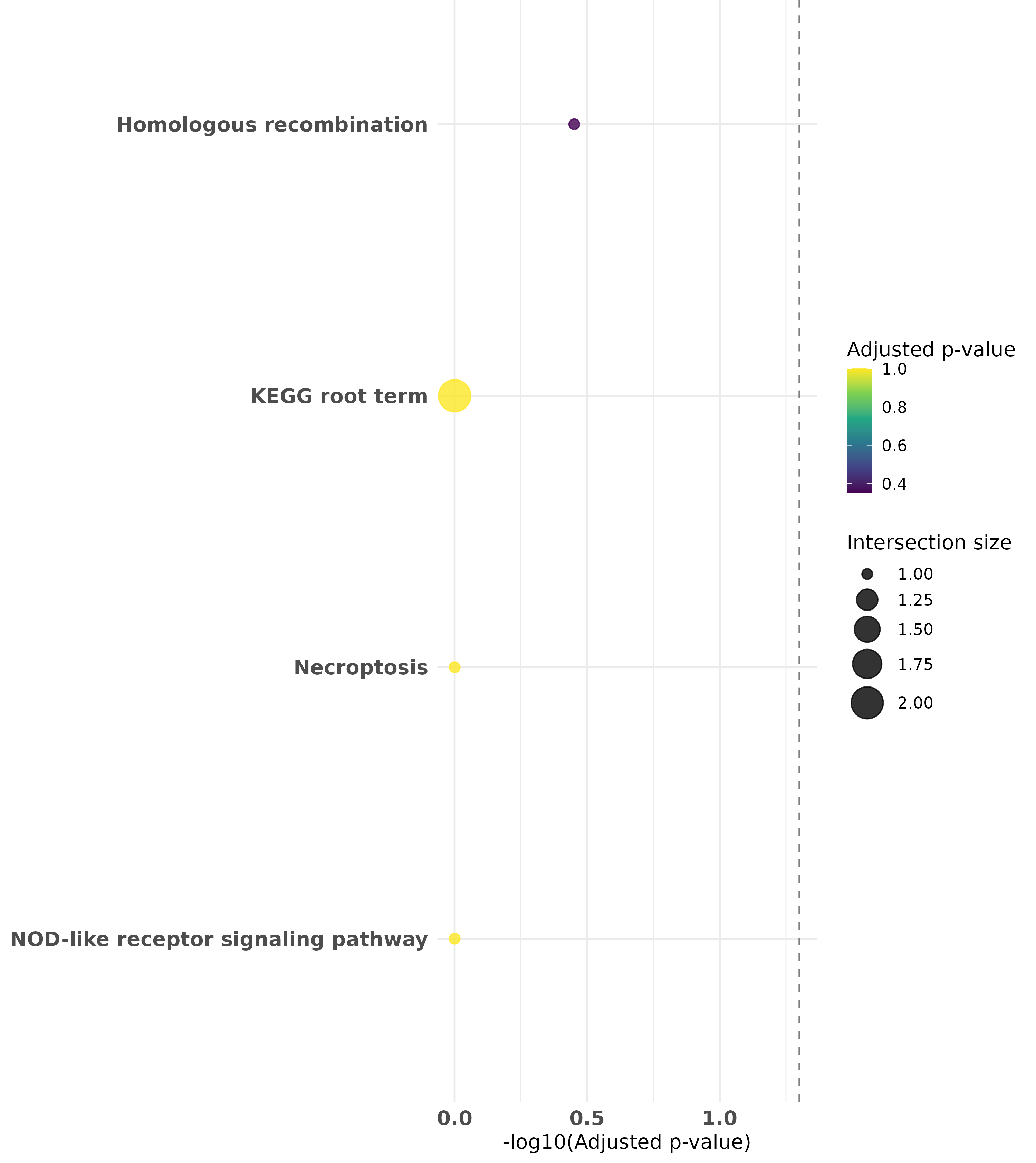

**D**

**C**

**Figure S2: Top enriched gene ontologies in Zinc-finger associated genes in cluster 4.** (**A**) Top enriched Biological processes (**B**) Top enriched Cellular Component (**C**) Top enriched Molecular Function (**D**) Top enriched KEGG pathways. Gene ontology enrichment was done with gProfiler using all annotated genes from *Danio rerio.* The dashed vertical line corresponds to statistical significance threshold (p-value of 0.05) on the x-axis.

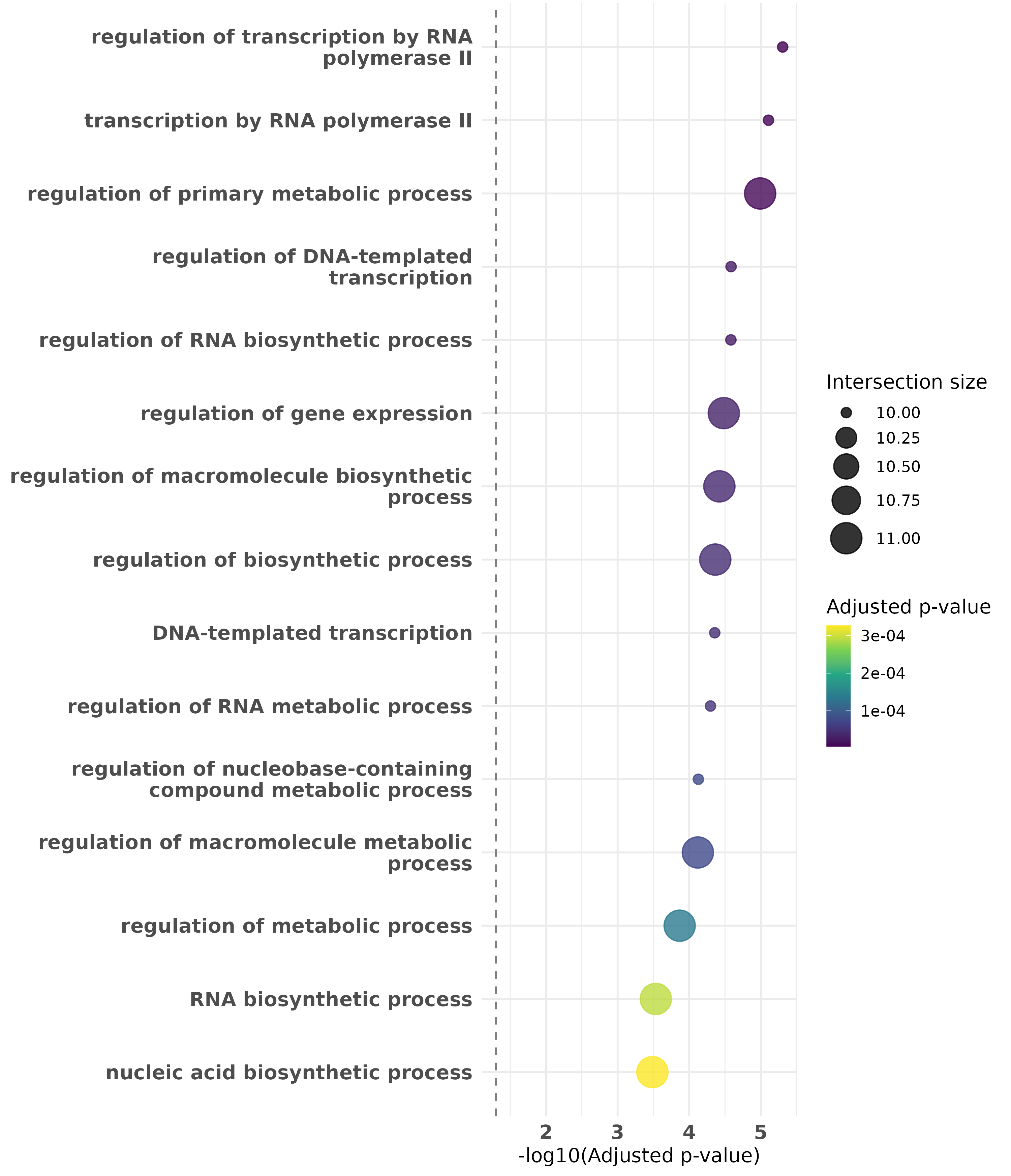

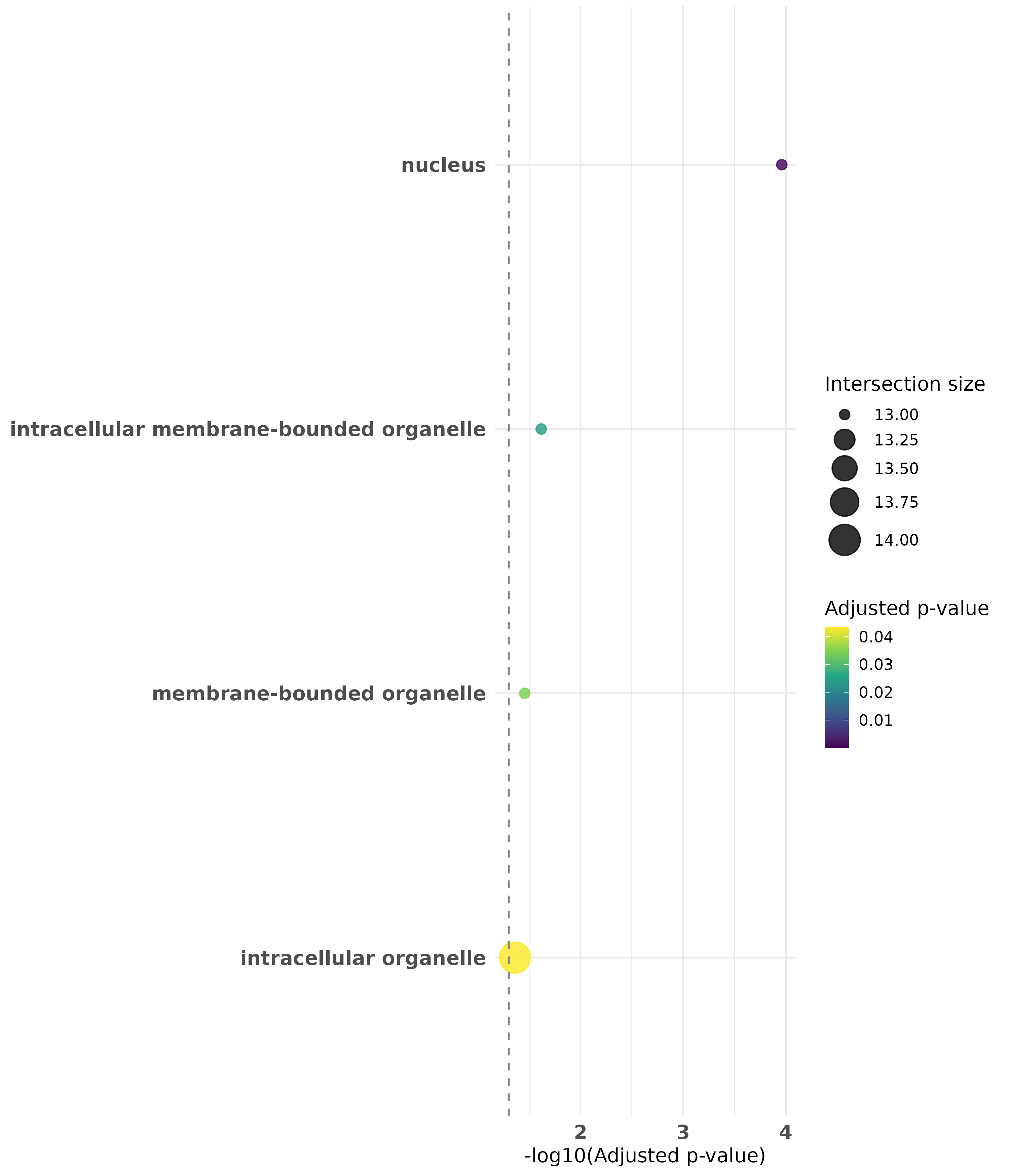

**B**

**A**

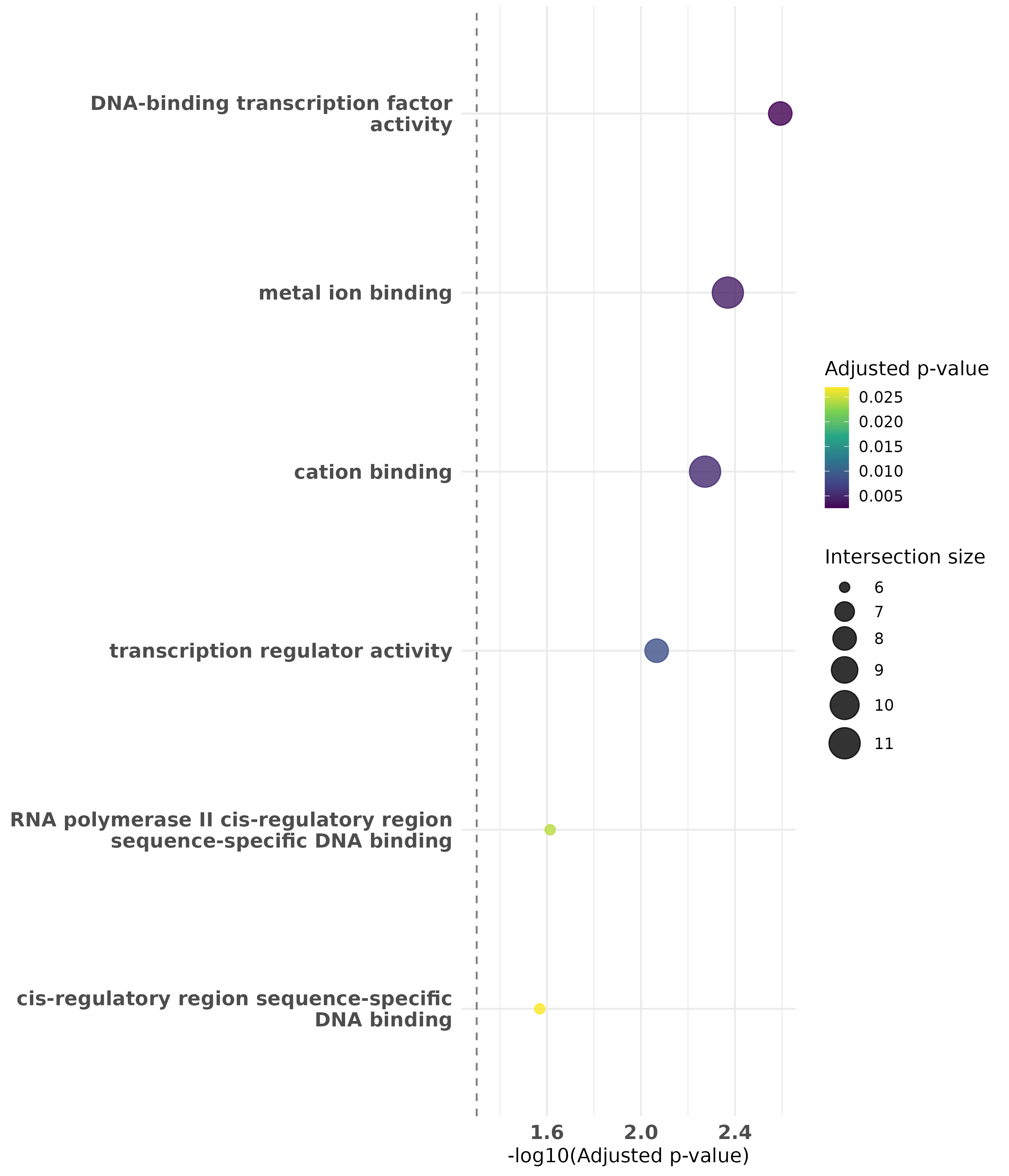

**C**

**Figure S3: Top enriched gene ontologies in Zinc-finger associated genes in cluster 5.** (**A**) Top enriched Biological processes (**B**) Top enriched Cellular Component (**C**) Top enriched Molecular Function. There are no significantly enriched KEGG pathways in this cluster. Gene ontology enrichment was done with gProfiler using all annotated genes from *Danio rerio.* The dashed vertical line corresponds to statistical significance threshold (p-value of 0.05) on the x-axis.

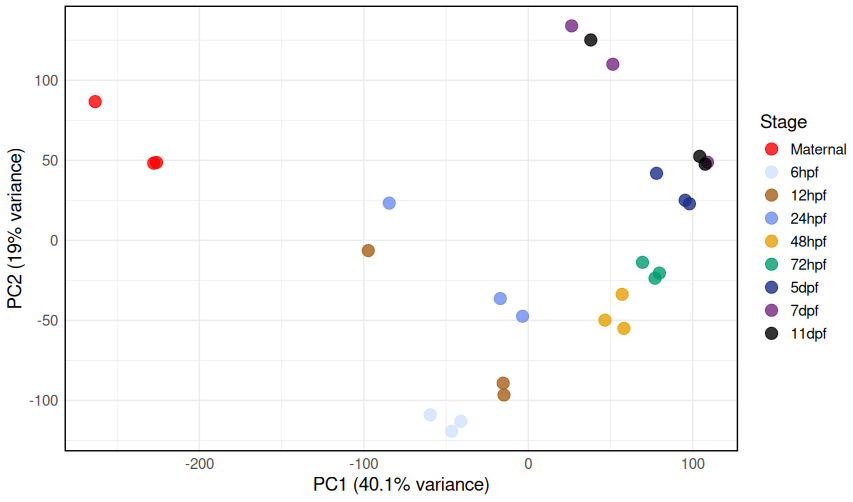

**A.**

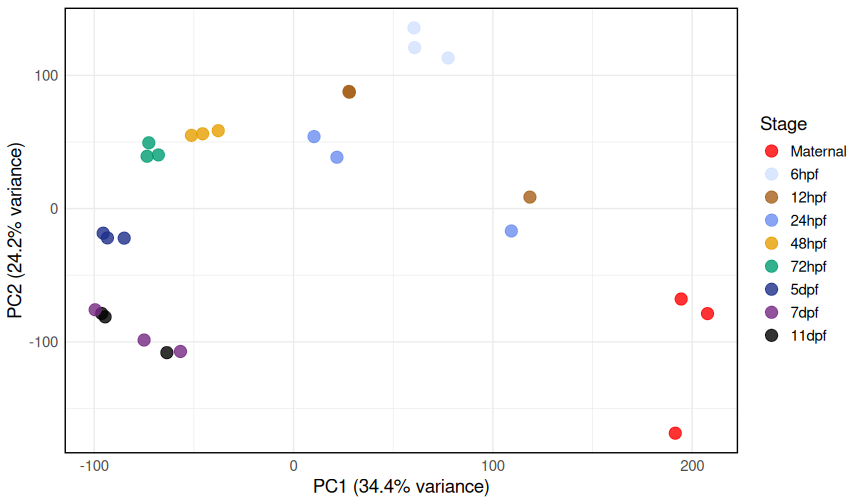

**B.**

**Figure S4:** **Principal component analysis of DESeq2 normalisation versus TPM across developmental timepoints.** (**A**). PCA of DESeq2 transformed size factor normalised counts. (**B**). PCA of TPM data.
